# Repurposing base editors for targeted knock-in and simultaneous knockouts to generate multiplex-edited allogeneic CAR T cells with minimal translocations

**DOI:** 10.1101/2025.09.15.676172

**Authors:** Viktor Glaser, Lily Jo Becker, Carla Fuster-García, Luis Huth, Ana-Maria Nitulescu, Yaolin Pu, Isabell Kassing, Laura Marie Hartmann, Christian Luca Flugel, Roberts Karklins, Shona Shaji, Marie Pouzolles, Maik Stein, Geoffroy Andrieux, Toni Cathomen, Hans-Dieter Volk, Petra Reinke, Jonas Kath, Dimitrios Laurin Wagner

## Abstract

The CRISPR-Cas system enables precise genome engineering of cell therapies. For allogeneic applications, multiplex editing is frequently required to improve efficacy, persistence, and safety. However, strategies involving multiple DNA double-strand breaks (DSBs) induce genotoxicity by provoking chromosomal aberrations. Base editors, which enable sequence changes without generating DSBs, are widely used for gene disruption, but their capacity for gene insertion remains unexplored. Here, we developed **B**ase **e**ditor-mediated **k**nock-**i**n (**BEKI**), a non-viral platform that allows targeted transgene insertion in parallel with multiplex gene disruption using a single base editor. Repurposing the Cas9 nickase domain of base editors generates paired nicks, inducing homology-directed repair (HDR). In human T cells, optimized guide RNA orientation and nick distance, together with HDR-enhancing modulators, enabled efficient transgene knock-in at the *TRAC*, *CD3ζ, B2M,* and *CD3ε* loci. Simultaneous base editing of multiple additional genes produced chimeric antigen receptor (CAR) T cells with increased cytokine secretion, drug resistance, and resistance to allo-rejection. Compared to multiplex editing with Cas9, BEKI markedly reduced chromosomal translocations. BEKI therefore provides a streamlined, scalable strategy for multiplex CAR T-cell engineering with a single enzyme, offering a safer route to clinical-grade manufacturing of off-the-shelf therapies for cancer and autoimmune diseases.

**Graphical Abstract:** 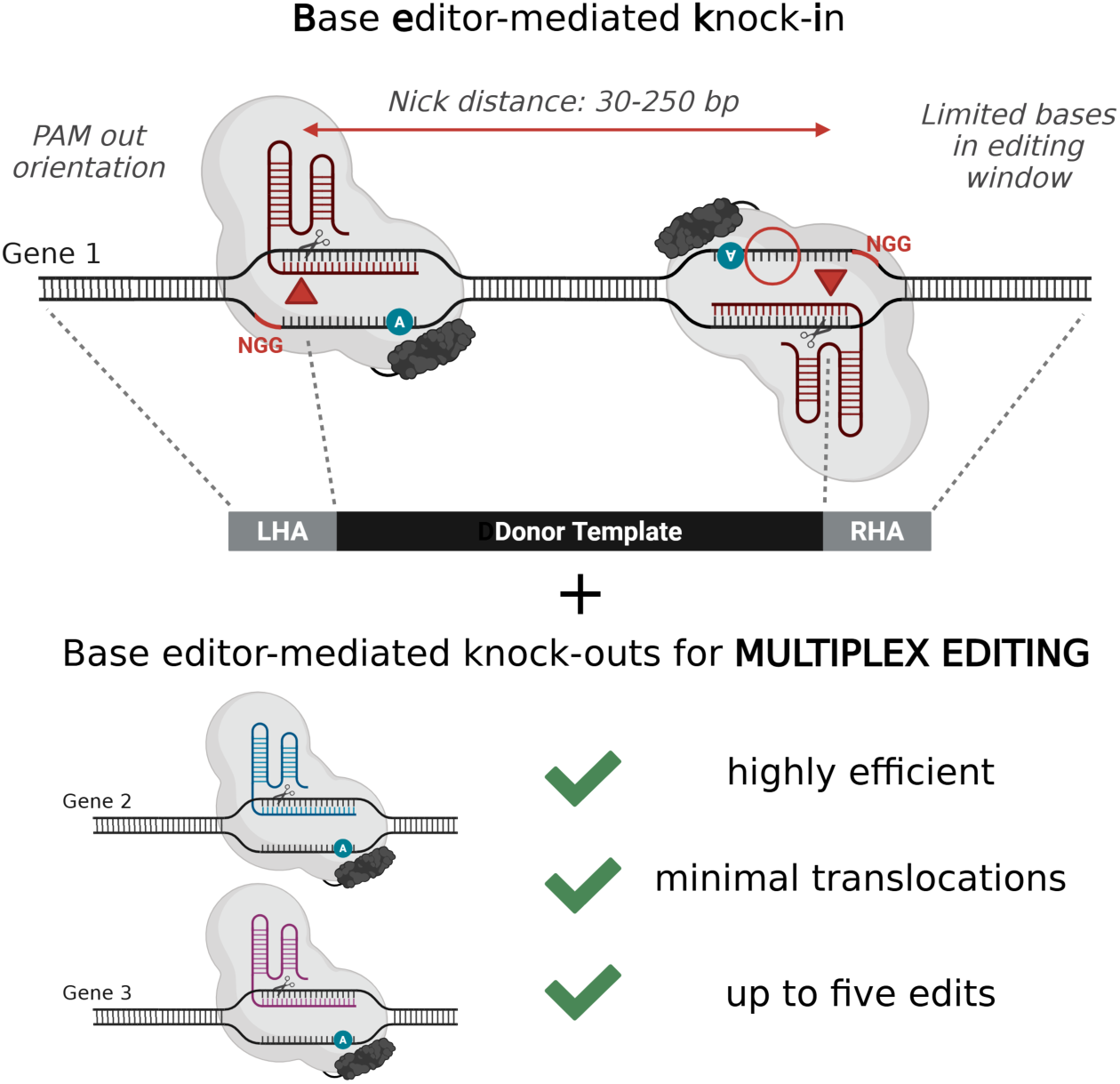

## Introduction

Recent advances in CRISPR-Cas-based gene editing have transformed cell and gene therapies by enabling programmable, site-specific modifications. Conventional Cas- endonucleases, such as *Streptococcus pyogenes* Cas9 (SpCas9) and *Acidaminococcus sp.* (AsCas12a), introduce DNA double-strand breaks (DSBs), which are repaired by endogenous DNA repair mechanisms: The non-homologous end joining (NHEJ) pathway rapidly ligates the DNA ends and often introduces small insertions or deletions (indels), which can be used for targeted gene knockout (KO). In contrast, homology-directed repair (HDR) leverages a homologous DNA template to achieve precise repair. By supplying an exogenous template, HDR can be harnessed for targeted transgene integration, commonly referred to as knock-in (KI). Leveraging these major DNA repair mechanisms enables precise engineering of cell therapies.

CRISPR-Cas-assisted HDR allows the non-viral generation of chimeric antigen receptor (CAR) T cells with controlled expression, for instance by integrating the CAR construct into defined genomic loci such as the *T cell receptor alpha constant* (*TRAC*) or *CD3*ζ gene, thereby eliminating the need for exogenous promoters^1–4^. Compared to virally transduced CAR T cells, characterized by random integration and exogenous promoter–driven expression, *TRAC*- and *CD3*ζ-integrated CAR T cells display reduced exhaustion and improved functionality^3,5^. Moreover, CRISPR KI at these loci disrupts the T cell receptor (TCR) complex, preventing graft-versus-host disease (GvHD) and forming the foundation of allogeneic CAR T cell therapies^6,7^. While non-virally engineered allogeneic CAR T cells promise scalable, off-the-shelf therapies derived from healthy donors at reduced cost^8^, they must be further engineered to evade rejection by the host’s immune system. This can be achieved by disrupting the genes *beta-2-microglobulin* (*B2M)* and *class II major histocompatibility complex transactivator* (*CIITA*) to abrogate Human Leukocyte Antigen (HLA) class I and II expression, respectively, and by introducing inhibitory receptors such as HLA-E to suppress NK cell-mediated rejection^9–11^.

Allogeneic CAR T cells may benefit from additional gene edits (such as KOs), including previously described strategies to prevent T cell exhaustion^12^ and improve function within the immunosuppressive tumor microenvironment^13^. Gene editing can also render CAR T cells resistant to immunosuppressive drugs. For example, KO of *FK506- binding protein 12* (*FKBP12*) confers resistance to tacrolimus and rapamycin^14^, and may thus enable treatment of patients who require systemic immunosuppression. Beyond these rational KO approaches, genome-wide CRISPR KO screens have been employed to uncover genes whose deletion can overcome CAR T cell dysfunction and improve long-term functionality^15–20^. One of the hits, KO of *RAS p21 protein activator 2* (*RASA2*), exhibited enhanced persistence and effector functions^21^. Although promising on their own, these edits gain full relevance in combination, especially for allogeneic CAR T cells, where multiplex engineering is likely essential for durable responses.

During multiplex editing with conventional Cas endonucleases, the simultaneous induction of multiple DSBs carries a high risk of genotoxicity, giving rise to structural variants (SVs) such as large deletions^22^, chromosome truncations^23^, chromosome loss^24^ and translocations, at both on-target and off-target sites^25^, potentially causing unintended gene fusions. While specific events from multiplex editing have not been linked to oncogenesis, certain recurrent translocations and fusions are established drivers of oncogenesis^26^. Given these concerns, mitigating genotoxicity is a critical consideration in the development of next-generation cellular therapies. Adenine base editors (ABEs) and cytosine base editors (CBEs) offer an alternative by employing a SpCas9 nickase (nCas9) fused to a deaminase to introduce predictable base substitutions without generating DSBs^27,28^. CBEs can introduce premature stop codons, while both ABEs and CBEs can disrupt splice sites to achieve efficient gene KO^29–31^.

Here, we introduce **b**ase **e**ditor-mediated **k**nock-**i**n (BEKI), a strategy that repurposes the nCas9-domain of base editors for double nick-mediated HDR, while enabling multiplex-editing with minimal translocations. We explored BEKI in four loci relevant to T cell engineering – *TRAC, CD3ζ, CD3ε*, and *B2M* – systematically assessing the impact of sgRNA design on editing efficiency. To further improve transgene insertion efficiency, we harnessed DNA repair modulation using small molecule inhibitors, previously used to enhance Cas9-based gene editing^32,33^. Utilizing the inherent base editing ability, BEKI enabled multiplex gene disruption by targeted splice site disruption. We demonstrated the generation of BEKI CAR T cells with simultaneous KOs of relevant genes for T cell engineering – *RASA2* and *FKBP12* – and assessed translocation frequencies using CAST-Seq and ddPCR. Finally, we generated BEKI CAR T cells with five edits introduced in parallel: *TRAC* KI of a CAR, and KOs of *B2M*, *CIITA*, *RASA2*, and *FKBP12*, conferring TCR replacement, immune evasion, enhanced cytokine production, and resistance to immunosuppressive drugs. This study establishes BEKI as a versatile platform for generating multiplex-engineered allogeneic CAR T cells by expanding the capability of base editors from precise single-base edits to simultaneous double nick-mediated insertion of therapeutic transgenes.

## Results

### Repurposing the nCas9 domain of base editors enables site-specific double nick-mediated integration of CAR transgenes

Previous studies using D10A nCas9 for double nicking have described that PAM orientation and the distance between nicks critically influence both HDR and NHEJ frequencies^34–36^. To develop an efficient double nick-mediated KI strategy, we designed 12 single-guide RNAs (sgRNAs) targeting the *TRAC* locus for in-frame insertion of a second generation CD19-specific CAR transgene (**Fig. 1A**). Specifically, 6 sgRNAs targeting the sense strand were combined with 6 sgRNAs targeting the antisense strand, creating a matrix of 36 distinct sgRNA combinations. The resulting sgRNA pairs varied in distance between the nick sites and exhibited either PAM-in or PAM-out orientation (**Fig. 1B)**. We electroporated activated primary human T cells with mRNA encoding a D10A nickase SpCas9^37^, two synthetic sgRNAs targeting opposed DNA strands and a linear double-stranded HDR donor template containing homology arms matching the genomic regions flanking the nick sites^36^. Among all sgRNA combinations tested, T1+T7 yielded the highest efficiency, with a mean of 27.5% CAR⁺ cells as measured by flow cytometry (**Fig. 1C)**. Notably, efficient editing outcomes with >5% CAR^+^ cells were only achieved using sgRNA combinations with a PAM-out orientation and a minimum distance of 21 bp between nicks (**Fig. 1C)**, aligning with previous nCas9-mediated KI studies in cell lines^34,35^. To test the proposed concept of BEKI, we replaced the nCas9 mRNA with ABE mRNA, achieving CAR integration rates of up to 16.0% (T1+T9) (**Fig. 1D**). Utilizing CBE for BEKI of a CAR yielded up to 9.0% of CAR+ cells (T2+T8) (**Fig. 1E**). While base editors can be repurposed for efficient double nick-mediated transgene insertion, fewer sgRNA combinations resulted in efficient KI compared to editing with nCas9 (**Fig. 1C-E**). Interestingly, we observed relatively higher CD3 KO efficiencies with BEKI compared to nCas9 editing in T cells transfected with sgRNA T4 targeting a splice site (**Suppl. Fig. 1)**. sgRNA pairs that enable robust nCas9 activity do not necessarily yield high BEKI efficiency (**Fig.1 C-E, Suppl. Fig. 2**). Further, BEKI using ABE or CBE generally resulted in lower KI- efficiencies compared to nCas9, while we did not observe a clear impact of the number of target nucleotides in the base editing window on BEKI rates (**Suppl. Fig. 3**).

**Fig 1.**
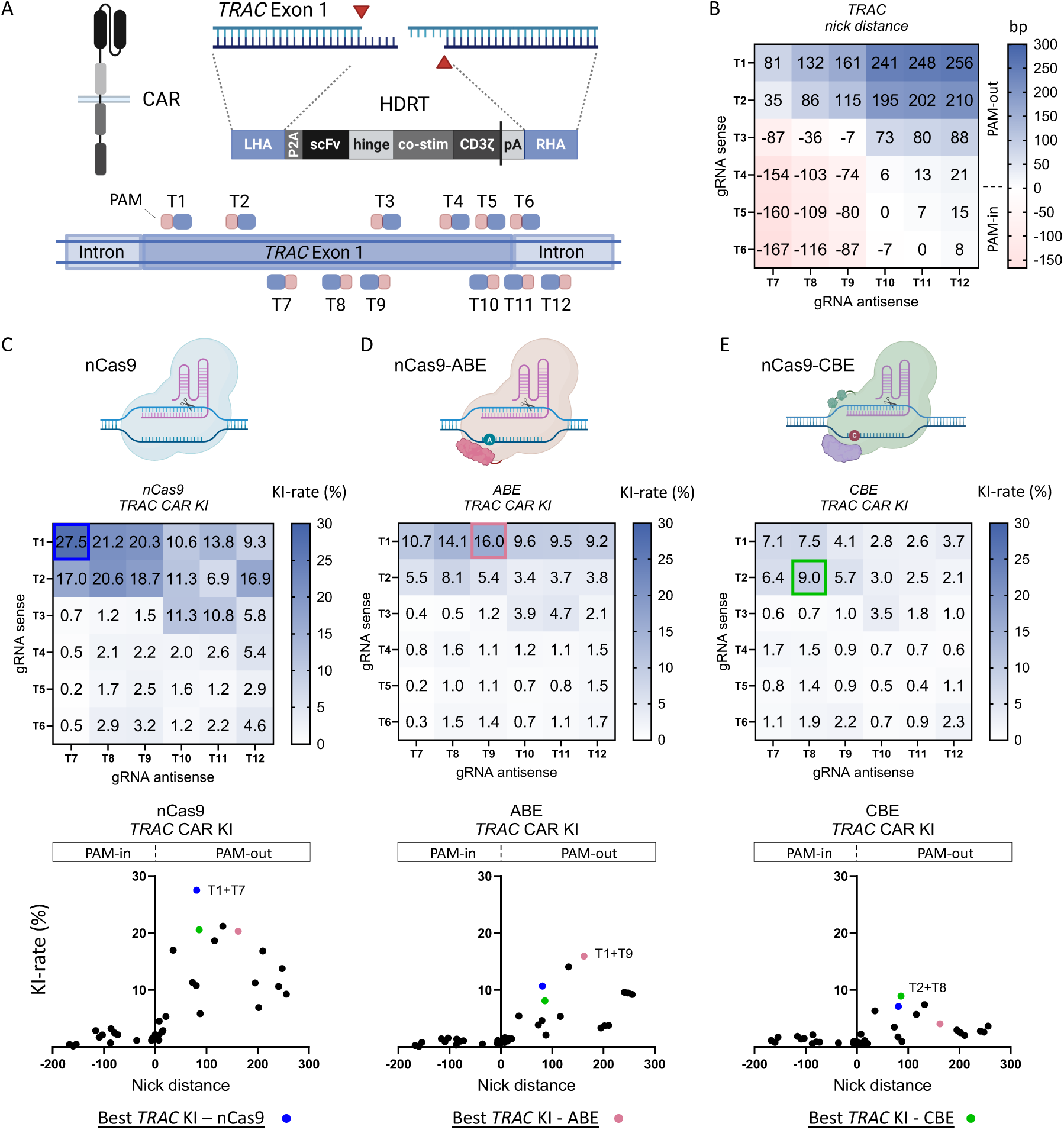
sgRNA pair design for site-specific integration of CAR transgenes using a double nick approach with nCas9 or base editors (BEKI). (A) Schematic overview of the double nick strategy for the insertion of a second generation CD19 CAR into the *TRAC* locus and the location and direction of the 12 sgRNAs used. (B) Nick distance of the 36 sgRNA combinations targeting opposed DNA strands. Nick distances of PAM-in oriented sgRNA pairs are displayed as negative values. (C) KI-efficiency using a D10A nCas9 was evaluated on day 4 after electroporation via flow cytometry (n=2 healthy donors). The results are displayed as a heatmap corresponding to B) and as a graph plotting the KI rate against the nick distance. (D) KI-efficiency using ABE8.20-m was evaluated on day 4 after electroporation via flow cytometry (n=2 healthy donors). The results are displayed as a heatmap corresponding to B) and as a graph plotting the KI rate against the nick distance. (E) KI-efficiency using a CBE (BE4) were evaluated on day 4 after electroporation via flow cytometry) (n=2 healthy donors). The results are displayed as a heatmap corresponding to B) and as a graph plotting the KI rate against the nick distance.

### BEKI enables efficient gene editing at multiple therapeutically relevant loci

To further validate BEKI using ABE, we tested three previously described KI strategies at loci relevant for T cell engineering: CD19-CAR insertion into *CD3ζ* exon 2 (utilizing a truncated transgene, as the genomic *CD3*ζ provides CAR signaling and transcript termination)^3^, HLA-E insertion into *B2M* exon 2 (to evade NK-mediated rejection)^11^, and CD19-scFv insertion into *CD3*ε exon 3 creating a T cell receptor fusion construct (TRuC)^38^ (**Fig. 2A-C**). Based on results from targeting the *TRAC* locus, we performed a less extensive sgRNA screening for these novel BEKI strategies, excluding sgRNA pairs with PAM-in orientation. BEKI edited cells were analyzed by flow cytometry to detect expression of the transgenic receptors (**Suppl. Fig. 4**). We achieved relatively high KI-efficiencies across all three loci, especially using the sgRNA combinations Z4+Z6/Z7 (*CD3ζ*, 13.5%/12.4%), B4+B6 (*B2M*, 9.2%) and E1+E4 (*CD3*ε, 10.3%) (**Fig. 2A-C**). Certain sgRNAs such as B4 targeting a splice sites in the *B2M* gene (published by Gaudelli et al.^39^) resulted in highly efficient KO (**Suppl. Fig. 5A**). The sgRNA pairs that yielded high KI-rates generally also induced efficient gene disruption at the target loci with KO rates of 61%, 77%, 67% and 48% for the *TRAC, CD3ζ, B2M* and *CD3*ε loci, respectively (T1+T9, Z4+Z7, B2+B6, E3 + E4; **Suppl. Fig. 5B**). For *B2M*, the combination of B4+B6 was selected for following experiments, considering both high HLA-E KI and *B2M* KO frequencies (**Suppl. Fig. 5B**). In general, we observed a trend of decreasing KI rates with increasing distances between the nicks, with the most efficient sgRNA combinations falling within an optimal editing window of 25-175 bp (**Fig. 2D**). Interestingly, the presence of adenines within the base editing window defined by the sgRNAs used did not appear to markedly affect KI frequencies (**Fig. 2E**- **F**). To sum up, efficient BEKI strategies could be established at all tested loci; and we conclude that optimal design requires sgRNA pairs with PAM-out orientation and nick distances between ∼25 to 175 bp.

**Fig 2.**
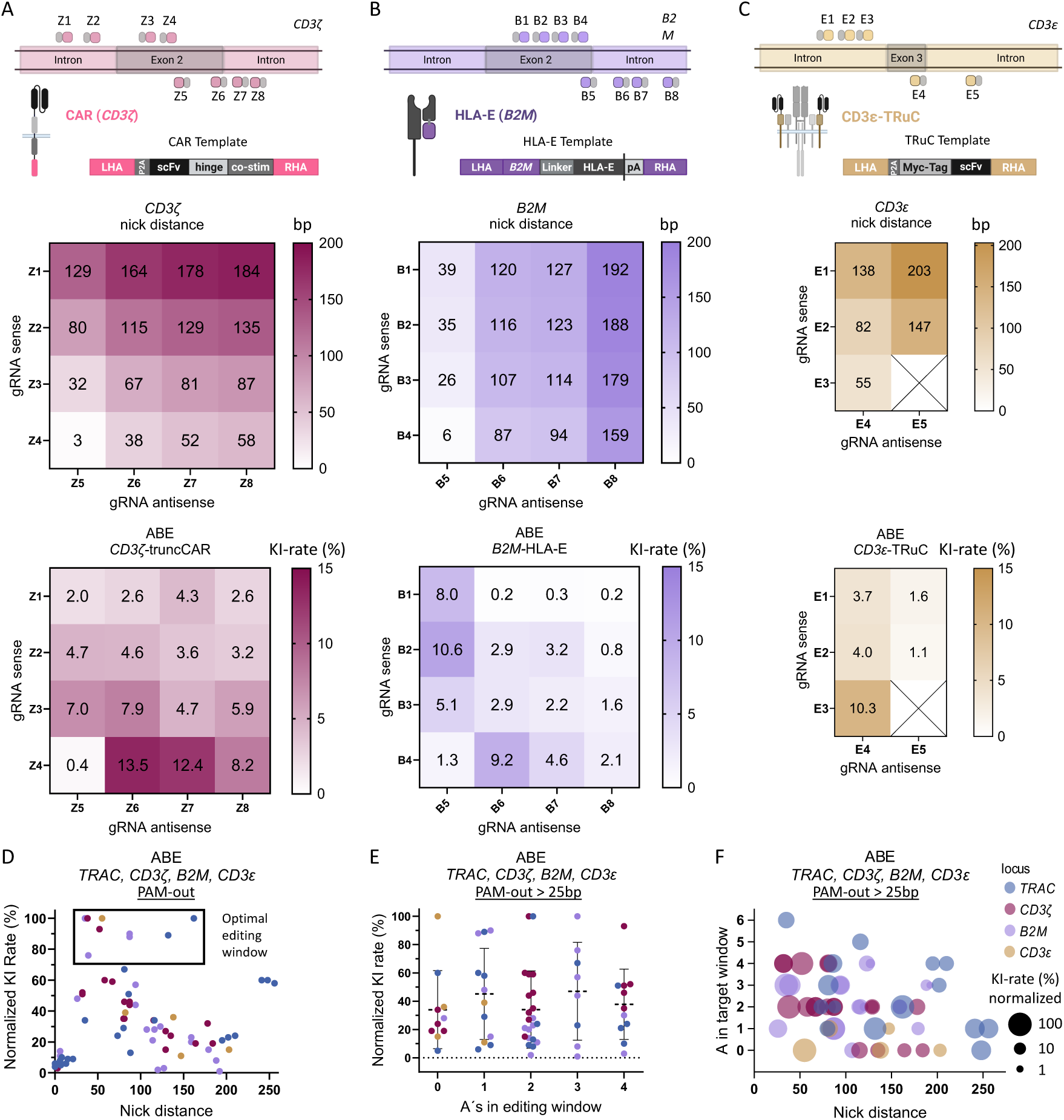
BEKI enables efficient gene editing at multiple therapeutically relevant loci. (**A**) Schematic overview of the double nick strategy for the insertion of a truncated (trunc)CD19 CAR into the *CD3*ζ locus and the location and direction of the 8 sgRNAs evaluated. Heatmaps are showing the nick distance of the 16 sgRNA combinations targeting opposed DNA strands and the KI-efficiency using ABE8.20-m, evaluated on day 4 after electroporation via flow cytometry (n=4 healthy donors). (**B**) The same was displayed for the KI of HLA-E into the *B2M* locus (8 sgRNAs, n=2) and (**C**) a CD19-specific TRuC into *CD3*ε (5 sgRNAs, n=3 healthy donors). (**D**) The KI-rate was normalized to the most efficient KI for each of the 4 tested loci (color) and plotted against the nick distance. The window for optimal editing was manually added. (**E**) The normalized KI-rate of editing at the 4 tested loci (color) was plotted against the number of adenines (A) in the editing window (5 and 6 As (n=1) are not shown) (**F**) The normalized KI-rate (circle size) of 4 tested loci (color) was plotted against the number of adenines (A) in the editing window and the nick distance.

### HDR enhancers improve BEKI efficiency and enable potent in vivo activity of CD19-CAR T cells

We and others have shown that selective inhibition of competing DNA repair pathways promotes HDR, thereby enhancing insertion of therapeutically relevant transgenes^4,40^. Based on this, we tested various NHEJ inhibitors for their effect on BEKI-mediated CAR insertion into *CD3ζ*, with the DNA-PK inhibitors Nedisertib (M3814) and AZD7648 showing the strongest enhancement (**Suppl. Fig. 6A**). Combining AZD7648 with Polθ inhibitors further improved insertion efficiencies across all tested sgRNA combinations compared to untreated controls, achieving KI frequencies of up to 40.2% (Z4+Z7, **Fig. 3A**). DNA-PK and Polθ inhibition also effectively improved CBE-mediated CAR insertion (**Fig. 3B).** Using the most effective sgRNA combinations at the *TRAC*, *CD3ζ*, *B2M*, and *CD3ε* loci, DNA-PK and Polθ inhibitor treatment resulted in an average 2.7- fold increase in ABE-mediated KI efficiency across all four targets (**Suppl. Fig. 6B**). We then evaluated the antitumor efficacy of BEKI-engineered *CD3ζ*-CAR T cells generated with DNA-PK and Polθ inhibitors in a xenograft model of acute lymphoblastic leukemia. Immunodeficient mice received 0.5 × 10⁶ luciferase-labeled CD19⁺ Nalm-6 cells intravenously, followed 7 days later by infusion of 0.5 × 10⁶ CAR⁺ cryopreserved T cells (**Fig. 3C**). *In vivo* bioluminescence imaging showed significantly reduced tumor growth in mice treated with *CD3ζ*-CAR T cells compared with mice receiving TCR/CD3-negative (*TRAC* KO) control T cells. (**Fig. 3C**). Consistently, 28 days post leukemia inoculation, leukemia cells were undetectable in blood from *CD3ζ*- CAR-treated mice (0/5) but detectable in 4/5 controls (**Fig. 3C**).

**Fig 3.**
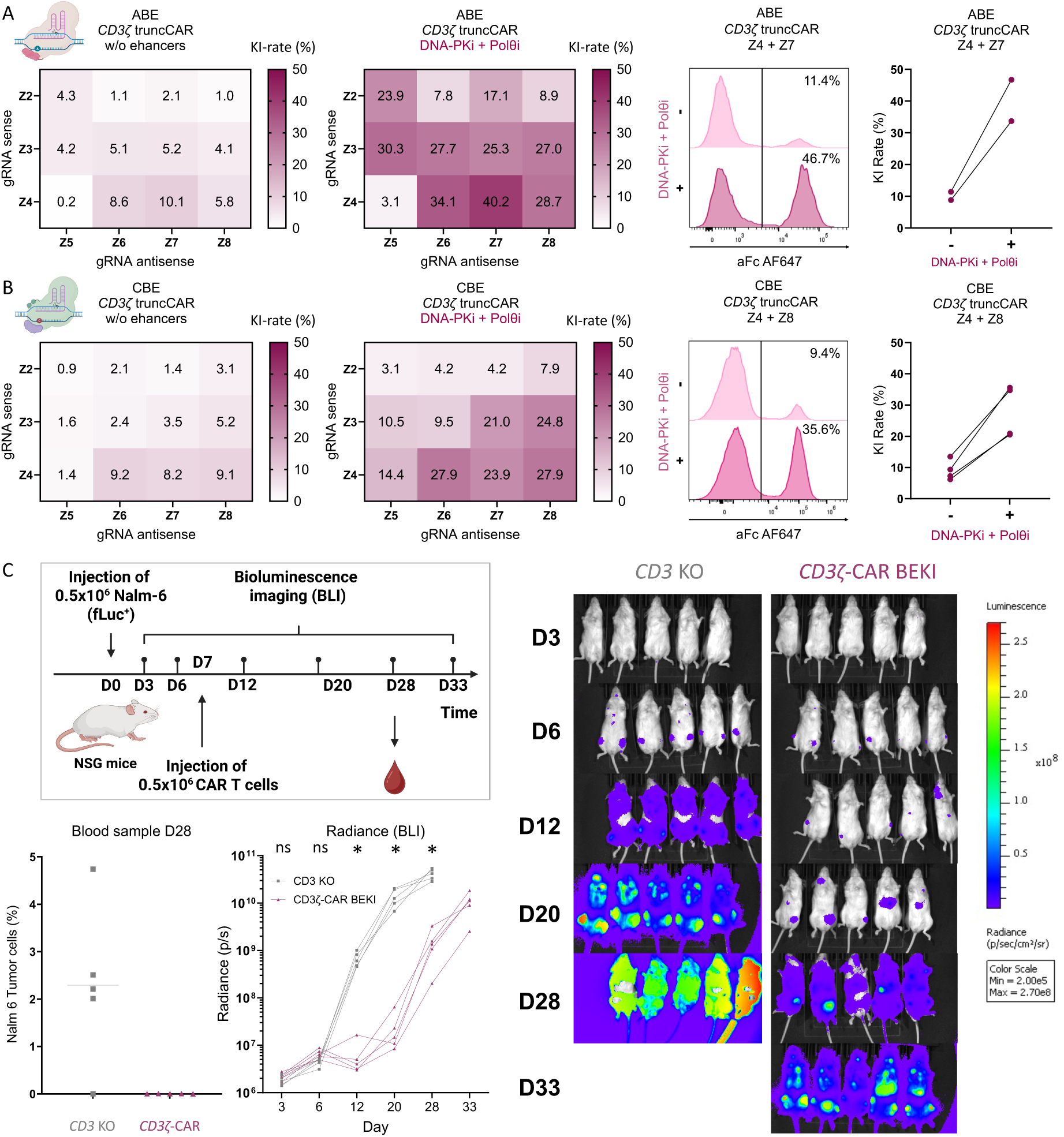
HDR enhancers improve BEKI editing efficiency and enable potent in vivo activity of CD19 CAR T cells. (**A**) ABE-mediated KI-efficiency of a truncCD19 CAR into *CD3*ζ without HDR enhancers. Z1 was excluded for this experiment due to low initial KI-rates (n=2 healthy donors). ABE-mediated KI-efficiency of a truncCD19 CAR into *CD3*ζ with DNA PK and Polθ inhibitors AZD7648 and Novobiocin (n=2 healthy donors). Representative flow cytometry histograms show different editing outcomes when adding DNA PK and Polθ inhibitors. KI-rate of the most efficient combination for *CD3*ζ KI using ABE: Z4+Z7 (n=2 healthy donors). (**B**) CBE-mediated KI-efficiency of a truncCD19 CAR into *CD3*ζ without HDR enhancers (n=2-6 healthy donors). CBE-mediated KI-efficiency of a truncCD19 CAR into *CD3*ζ with DNA PK and Polθ inhibitors AZD7648 and ART558 (n=2-6 healthy donors). Representative flow cytometry histograms show different editing outcomes when adding DNA PK and Polθ inhibitors. KI-rate of the most efficient combination for *CD3*ζ KI using CBE: Z4+Z8 (n=4 healthy donors). (**C**) Schematic overview of an acute lymphoblastic leukemia xenograft mouse model using luciferase-labeled Nalm-6 (CD19+) tumor cells. 7 days post Nalm-6 administration, 0.5×10^6^ cryopreserved CAR+ T cells were injected systemically. A blood sample was taken from all mice on day 28 after injection of the Nalm-6 tumor cells. Flow cytometric analysis was performed to quantify the frequency of GFP+ Nalm6 cells. Additionally, tumor burden was assessed via bioluminescence imaging (n = 5 mice). A multiple unpaired T test using the two-stage step-up method (Benjamini, Krieger, and Yekutieli) of log-transformed bioluminescence imaging data Asterisks represent the following p-values: (ns: p ≥ 0.05; *: p < 0.05; ^∗∗^ : p < 0.01; ^∗∗∗^ : p < 0.001).

### Multiplex base editing enhances BEKI CAR T cell function in the presence of immunosuppressants and enables allogeneic application

Leveraging the inherent base editing capability of the BEKI platform for simultaneous transgene KI and multiple base edits, we aimed to enhance *TRAC*-CAR T cells by introducing additional KOs of *RASA2*^21^ and *FKBP12*^14^. ABE-mediated splice site disruption of *FKBP12* conferred resistance to the calcineurin inhibitor tacrolimus, as indicated by sustained cytokine production in its presence (**Suppl. Fig. 7A**). Efficient splice site disruption of *RASA2* enhanced cytokine production upon polyclonal activation (**Suppl. Fig. 7B**). Although *RASA2* KO has previously been linked to enhanced proliferation and cytotoxicity in immunosuppressive environments, such as during tacrolimus treatment^21^, *RASA2* base editing alone did not sufficiently restore cytokine production under tacrolimus exposure in our hands (**Suppl. Fig. 7C**). Consequently, we compared two strategies for combining CAR KI with disruption of both ***<u>R</u>****ASA2* and ***<u>F</u>****KBP12* (CAR + *RF* KO): (1) BEKI, in which the ABE mediated both CAR KI and the KOs, and (2) a control approach with Cas12a-mediated CAR KI with ABE-mediated KOs. Both strategies successfully enabled tumor-specific cytokine production in the presence of tacrolimus (**Suppl. Fig. 7C**). Next, we successfully generated quintuple-edited BEKI CAR T cells combining ***<u>T</u>****RAC*-CAR KI with base editor-mediated KOs of ***<u>B</u>****2M*, ***<u>C</u>****IITA, **<u>R</u>**ASA2* and ***<u>F</u>****KBP12* (CAR+*TBCRF* KO) (**Suppl. Fig. 7D**). Importantly, the addition of two or four sgRNAs for splice site disruption did not compromise CAR KI rates, while reaching high KO efficiencies for HLA class I and II (**Suppl. Fig. 7E**). These quintuple-edited BEKI CAR T cells evaded allo-specific T cell cytotoxicity (**Suppl. Fig. 7F**) and maintained CAR function under therapeutic rapamycin concentrations (**Suppl. Fig. 7G**), supporting their use in allogeneic settings and in patients receiving immunosuppressive therapy.

### BEKI minimizes chromosomal translocations during simultaneous KI and multi- gene KO

Aiming to benchmark translocation risk, we compared BEKI against two other multiplex strategies for generating ***<u>T</u>****RAC*-CAR T cells in combination with ***<u>R</u>****ASA2* and ***<u>F</u>****KBP12* KOs (CAR + TRF KO): (1) Cas9 for CAR KI and the additional KOs, expected to induce translocations between the on-target DSBs, and (2) Cas12a for CAR-KI plus ABE for the KOs, previously shown not to induce translocations^41^. As a positive control for inducing translocations, we included Cas9-mediated triple KO without an HDR template (KO only) (**Fig. 4A**). Cas12a + ABE achieved the highest CAR integration frequencies, whereas BEKI matched Cas9 in KI efficiency but with lower CD3 KO frequencies; *RASA2* and *FKBP12* KOs, however, were most efficient with ABE (**Fig. 4B**). To avoid interference with the HDR template, we assessed genomic integrity by CAST-Seq with the KO-only samples (**Fig. 4C**). These KO-only samples showed similar KO frequency trends as when edited with DNA template (**Suppl. Fig. 8A**). CAST-Seq identified several types of genomic alterations: large chromosomal aberrations at the *TRAC* target site (**Suppl. Fig. 8B**), translocations between *TRAC* and the two other on-target loci (*FKBP12* and *RASA2*), as well as off-target–mediated translocations (OMTs) (**Fig. 4C**). Chromosomal rearrangements between *TRAC*, *RASA2*, and *FKBP12* were detected with all strategies (**Fig. 4C**). Furthermore, while OMTs were frequent with Cas9 multiplex editing, only one was detected with Cas12a + ABE, and none with ABE (BEKI) (**Fig. 4C**). We next quantified translocation frequencies by ddPCR, designing probes for all six possible balanced translocations between *TRAC*, *RASA2*, and *FKBP12*. These translocations were readily detected in Cas9-edited cells ((I) *TRF* KO, (II) CAR + *TRF* KO), with frequencies up to 4%, whereas such translocations were undetectable by ddPCR in cells edited using either (III) Cas12a + ABE or (IV) multiplex BEKI, demonstrating their superiority in maintaining genomic integrity (**Fig. 4D**).

**Fig 4.**
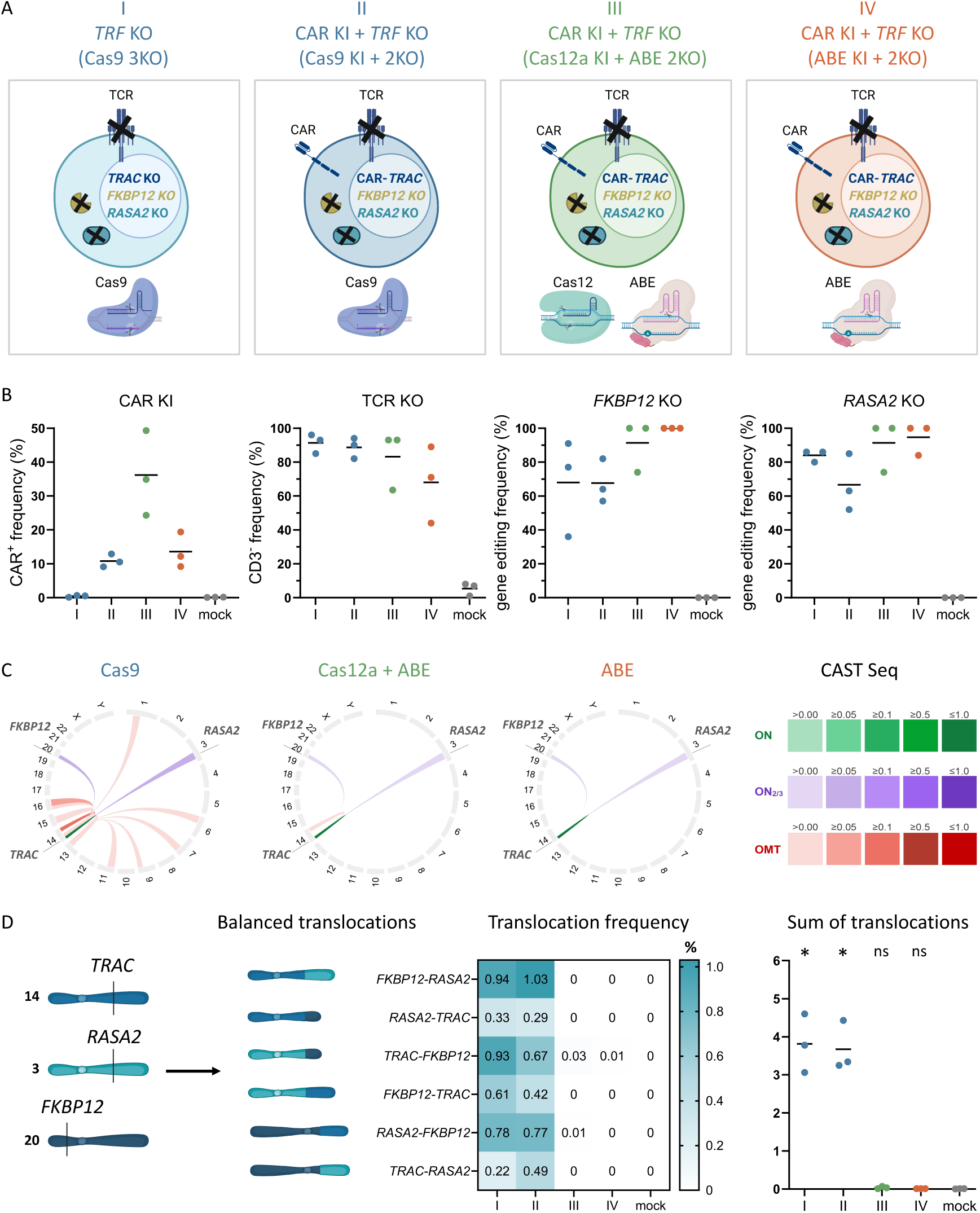
BEKI minimizes chromosomal translocations during simultaneous knock-in and multi-gene knockout. (**A**) Schematic overview of the different gene editing strategies targeting the *TRAC, RASA2* and *FKBP12* gene to create cells with (I) *TRF* KO using Cas9, (II) CAR KI + *TRF* KO using Cas9, (III) CAR KI + *TRF* KO using Cas12a for the KI and ABE for the *RF* KO and (IV) CAR KI + *TRF* KO using ABE (**B**) Gene editing efficiency of CAR KI and *TRAC* KO (CD3 staining) was determined by flow cytometry. KO- or BE-efficiency of *RASA2* and *FKBP12* gene were assessed using sanger sequencing followed by indel analysis using ICE CRISPR Analysis. 2025. v3.0. (EditCo Bio) or base editing frequency via EditR. (**C**) Circos plot shows results of chromosomal rearrangements detected in cells edited with Cas9, Cas12a + ABE or ABE using CAST-Seq (excluding an HDR donor template). On-target (ON) genomic aberrations are indicated in green (*TRAC*) or purple (*RASA2, FKBP12*), and off-target mediated rearrangement between the *TRAC* target site and off-target (OMT) site in red. The color intensities of the arcs directly reflect the relative frequency of translocations, normalized to the most frequent event, with the corresponding range indicated in the heatmap legend on the right. (**D**) Frequencies of balanced translocations between the on-target genes were determined by ddPCR. They are shown for all six individual translocations and as the sum of all translocations detected in mock, (I) TRF KO (Cas9) or CAR KI + TRF KO ((II) Cas9, (III) Cas12a+ABE, (IV) ABE) conditions (n = 3 healthy donors). Statistical analysis of ddPCR was performed using a one-way ANOVA of matched data with Geisser-Greenhouse correction. Multiple comparisons were performed by comparing the mean of each column with the mean of every other column and corrected by the Turkey test. Asterisks represent the following p-values: (ns: p ≥ 0.05; *: p < 0.05; ** : p < 0.01; *** : p < 0.001).

## Discussion

Here we show that base editors can be repurposed for targeted KIs. By exploiting their D10A nCas9 domain to induce double nicks and stimulate HDR, we extend their utility beyond single nucleotide editing. Coupling this KI capacity with their established ability to mediate gene KOs, BEKI enables complex engineering with a single enzyme. This approach effectively minimizes chromosomal translocations, a major drawback of conventional Cas9-based multiplex editing. To guide the design of new BEKI strategies, we note that, consistent with early nCas9-based double-nick approaches^34,35^, efficient transgene KI was achieved when sgRNAs were selected in a PAM-out orientation creating nicks on opposite DNA strands at a distance of 25– 175 bp. Potential base editing mediated through the sgRNA pair used for double nicking had no evident impact on BEKI efficiency, likely due to the PAM-distal position of the catalyzed base substitution, a region where sgRNA mismatches are well tolerated by Cas9^42^. However, we still recommend selecting sgRNAs with minimal editable bases within the activity window of the base editor.

Using BEKI at the *TRAC* locus, we observed KO rates exceeding 60% with sgRNA pairs spaced up to 250 bp apart (**Suppl. Fig. 1**), contrasting a previous report that increasing nick distance reduces NHEJ frequencies^36^. In the context of allogenic CAR T cell therapies, eliminating residual CD3^+^ cells is crucial to prevent GvHD and enables selective enrichment of CAR^+^ cells, particularly as the CD3 KO efficiency using BEKI is lower than typically achieved with Cas9^43^. To achieve high KO rates, sgRNAs designed for base editing of splice sites were effectively repurposed for KI strategies (*TRAC* sgRNA T4, **Suppl. Fig. 1,** *B2M* sgRNA B4 **Suppl. Fig. 5**). Surprisingly, the *CD3*ζ targeting sgRNA Z3 caused high KO rates (**Suppl. Fig. 5**), likely due to ABE induced amino acid changes (L34P/L35P) in the transmembrane domain of CD3ζ potentially disrupting the α-helix structure of the transmembrane domain and disrupting TCR assembly^44^. These findings highlight the importance of considering potential amino acid-altering base edits introduced by the sgRNA pair used for double nicking.

At the 4 targeted loci, baseline KI efficiencies with BEKI were modest, averaging around 10% (**Fig. 1-2**), underscoring the need for strategies to increase editing rates to achieve clinically relevant cell yields. Pharmacologic inhibition of DNA-PK has been shown to increase KI rates but at the cost of promoting large deletions, while Polθ inhibition can counteract this undesired effect^4,32,33^. In line with these findings, we applied the combination of DNA-PK and Polθ inhibitors during editing, which substantially improved BEKI-mediated CAR KI efficiencies (**Fig. 3**), although potential effects on genomic stability and cellular fitness warrant further investigation. A concurrent study (INSERT) similarly demonstrated that base editors can be leveraged for double-nick–mediated transgene insertion, but relied on AAV-delivered donor templates^45^. INSERT further improved KI efficiency through a ‘double-tap’ approach, in which additional sgRNAs target base-editing byproducts to enable iterative nicking. While AAV donor delivery may increase KI rates compared to dsDNA, viral templates introduce added complexity, cost, and regulatory burden^46,47^. Future improvements in KI rates could come from reducing donor toxicity, e.g. via transient DNA sensing inhibition^4^, or by using modified single-stranded DNA (ssDNA) donor templates^48,49^, which would allow higher template doses and promote HDR, although ssDNA may be prone to deamination by base editors^50^.

*CD3ζ*-integrated CD19-CAR T cells generated with BEKI under transient DNA-PK and Polθ inhibition demonstrated in vivo antitumor activity in a xenograft Nalm-6 lymphoma model (**Fig. 3C**). We then combined CAR KI with *RASA2* and *FKBP12* disruption, yielding triple-edited CAR T cells that displayed enhanced effector function under pharmacologic immunosuppression with tacrolimus or rapamycin. These findings illustrate the dual functionality of BEKI to mediate targeted KI and multiplex KO using a single enzyme (**Suppl. Fig. 7, Fig. 3A-B**). Similar to approaches that confer resistance to lymphodepleting agents^25,51^, engineering resistance to immunosuppressants could improve the persistence of allogeneic cells. We extended this strategy with additional *B2M* and *CIITA* edits for immune evasion, illustrating that BEKI can sustain progressively complex editing strategies. The resulting quintuple- edited allogeneic CAR T cells showed enhanced anti-tumor function and were protected from allo-specific rejection (**Suppl. Fig. 7**).

A key challenge in increasing the complexity of gene editing for cellular therapies is maintaining genomic integrity while keeping manufacturing feasible. Multiple nuclease-induced DSBs exert selective pressure that favors the survival of p53-deficient cells, whose reduced DNA damage sensitivity predisposes them to malignant transformation^52,53^. To mitigate the risk of DSB-induced genotoxicity and its consequences, several strategies have been explored. Viral gene transfer combined with multiplex base editing reduces DSB-associated genotoxicity, but requires multiple editing steps and complicates manufacturing^29,54,55^. Transposases combined with base editing enable non-viral multiplexed editing in a single transfection^56^. However, random transgene integration by viral vectors and transposases has been associated with insertional mutagenesis and cases of secondary malignancies^57,58^. Gene editing methods that achieve high HDR rates, either by using AAV-delivered donor templates^59^ or by selecting KI loci that inherently allow enrichment of edited cells ^60,61^, minimize translocation risk by favoring the intended editing outcome. We and others have described multiplex editing platforms that combine CRISPR-Cas nuclease-mediated KI and multiple KOs in a single step, using orthogonal enzymes^41^ or aptamer- extended sgRNAs^62^ to direct DSBs and base edits to distinct loci. While these approaches enable precise multiplex editing with reduced translocations risks, the complexity of delivering multiple components limits clinical translation. New strategies such as peptide-mediated ribonucleoprotein (RNP) delivery (PERC)^63^ and lipid nanoparticles (LNPs)^64^ may improve feasibility by lowering the toxicities of electroporation and DNA template delivery, and by enabling temporal separation of editing steps.

The BEKI platform overcomes many of these challenges by enabling precise, non-viral KI and multiplex gene KOs in a single step with a single enzyme. By restricting DSBs to the KI site and employing base editing for gene KOs, this approach minimizes translocation risks and simplifies manufacturing by relying on a single editor, eliminating the need for orthogonal enzymes or viral components. Harnessing base editing instead of Cas9-editing for multiplex engineering can improve T cell therapies by increasing both viability and editing rates^54,65^. Together, these features establish BEKI as a streamlined non-viral platform for the safe and efficient multiplex engineering in a single step, exemplified by the generation of quintuple-edited allogeneic CAR T cells, representing a step towards more effective and accessible T cell therapies.

## Material and Methods

### PBMC isolation and T cell enrichment

Peripheral blood was collected from healthy adult donors with informed consent under Charité ethics approval (EA4/091/19 and EA1/052/22), in accordance with the Declaration of Helsinki. Blood drawn into 10 mL lithium-heparin tubes (BD) was diluted 1:1 with sterile PBS (Gibco) and layered onto 15 mL of Pancoll (PAN Biotech) or Biocoll (Bio&SELL) in Leucosep™ tubes (Greiner). Samples were centrifuged at 800 × g for 20 minutes at room temperature without brake. The PBMC layer was collected from the interface, washed twice with PBS (300 × g, 10 min), and viable cells were counted using a Neubauer chamber and trypan blue exclusion (Sigma-Aldrich).

Prior to T cell seeding, 24-well tissue culture plates were coated with 1 µg/mL each of anti-CD3 (Invitrogen) and anti-CD28 (BioLegend) in 0.5 mL/well sterile water (Ampuwa), sealed with parafilm, and incubated at 4 °C for 24 h. Human CD3⁺ T cells were enriched from PBMCs via magnetic-activated cell sorting (MACS, Miltenyi Biotec) following the manufacturer’s protocol. In brief, cells were resuspended in cold MACS buffer (PBS + 2% heat-inactivated FCS + 2 mM EDTA), incubated with anti-CD3- conjugated magnetic beads for 15 min at 4 °C, washed, filtered (30 µm), and separated using LS columns (Miltenyi Biotec). Isolated cells were counted and seeded at 0.75 × 10⁶ cells/mL into pre-washed αCD3/αCD28-coated wells.

### Cell culture

T cells were cultured at 37 °C and 5% CO₂ in CTL medium consisting of 1:1 Advanced RPMI and Click’s Medium with 1% GlutaMAX supplemented with 10% heat-inactivated FCS (Biochrom or Sigma-Aldrich), 10 ng/mL IL-7, and 5 ng/mL IL-15 (CellGenix). Antibiotics were omitted due to their negative impact on post-electroporation viability. Nalm-6 cells were cultured in RPMI 1640 with 10% FCS, 100 IU/mL penicillin, and 100 μg/mL streptomycin, and passaged every 2–3 days.

### Plasmid construction and cloning

Cloning strategies were planned with SnapGene. Multiple-fragment In-Fusion cloning (Takara Bio) was performed using purified PCR fragments (KAPA HiFi HotStart, Roche) designed with primers generating 15 bp overlaps. In-Fusion reactions (5 μL) were transformed into 10 μL Stellar Competent *E. coli*, plated on LB-ampicillin agar, and incubated at 37 °C overnight. Colonies were screened by PCR using M13 universal primers flanking the pUC19 insertion site. Positive clones were grown in 5 mL LB/ampicillin overnight, and plasmids were purified (ZymoPURE Mini Prep Kit, Zymo Research). Sequence integrity of plasmids was confirmed via Sanger sequencing (LGC Genomics, Eurofins).

### In vitro transcription (IVT) of mRNA

ABE8.20-m (Addgene #136300,) and BE4_RrA3F (Addgene #138340) from Gaudelli et al.^39,66^ were cloned into plasmids containing a dT7 promoter, Kozak sequence, and 5′/3′ UTRs (TriLink). In addition, a D10A nickase Cas9 was derived from ABE8.20-m using In-Fusion cloning (Takara), and a wild-type Cas9 construct was generated by correcting the D10A mutation. Linear PCR templates were generated using KAPA HiFi HotStart (Roche), installing the correct T7 promoter sequence and a 120 bp polyA tail. PCR products were purified (Zymo DNA Clean & Concentrator-5) and IVT was performed using the HiScribe™ T7 High Yield RNA Synthesis Kit (NEB) with N1- methyl-pseudouridine (TriLink) and CleanCap AG (TriLink) for co-transcriptional capping. Template DNA was removed with DNase I (NEB), and mRNA was purified using the Monarch® RNA Cleanup Kit (NEB). RNA integrity was assessed via 1.5% denaturing agarose gel electrophoresis using RNA loading dye and ssRNA ladder (NEB). Finally, the mRNA concentration was measured using a Qubit fluorometer with the RNA BR Assay (Thermo Fisher Sientific), diluted to 2 μg/μL and stored at −80 °C.

### Generation and modification of dsDNA HDRTs for CAR insertion

HDR donor templates (HDRTs) were engineered for targeted insertion of a second-generation CD19 chimeric antigen receptor (CAR) into the *TRAC* locus, a truncated CAR lacking CD3ζ into the *CD3ζ* locus, an HLA-E–B2M fusion construct into *B2M* and a TCR fusion construct (TRuC) into *CD3ε*. Previously described HDRTs (*TRAC*^2^*, CD3ζ*^3^*, B2M*^11^*, CD3ε*^38^) were modified for improved compatibility with nickase-based editing by adjusting homology arms (HAs) to flank the generated DNA nicks (**Suppl. Table 1**). In some cases, HAs were adapted to accommodate multiple proximal sgRNAs by positioning HA ends within 35 bp of the nick site. Where HDRTs included sequences homologous to the sgRNA, silent PAM mutations were introduced to prevent re-cleavage. Multiple-fragment In-Fusion cloning (Clontech, Takara) was used to assemble HDRTs from overlapping PCR products generated with KAPA HiFi HotStart 2x Readymix (Roche). For HDRT generation, 500 μL PCR reactions were performed and purified using AMPure XP paramagnetic beads (Beckman Coulter) with two 70% ethanol washes. Purified HDRTs were quantified using a Nanodrop spectrophotometer (Thermo Fisher Sientific) and adjusted to a concentration of 1 or 2 μg/μL in nuclease-free water.

### Gene Editing

Primary human T cells were harvested ∼48 hours after anti-CD3/CD28 stimulation, counted, and washed twice in sterile PBS by centrifugation at 100 × g for 10 min at room temperature (RT).

#### mRNA-based knock-in

Double nick-mediated knock-ins were performed by co-electroporating: (1) HDR donor template (1 μL, 1–2 μg/μL), (2) two strand-specific sgRNAs (0.48 μL each, 100 μM; IDT), and (3) mRNA encoding the base or nuclease editor (1 μL, 2 μg/μL). Optionally, additional sgRNAs for base editing were included. Reagents were assembled in PCR strips by adding components sequentially. Synthetic, chemically modified sgRNAs (or crRNAs for Cas12a; IDT) were resuspended in nuclease-free 1×TE buffer and stored at −20 °C (Suppl. Table 2). sgRNAs for splice site disruption using base editors were designed using SpliceR^30^.

#### Cas12a Ribonucleoprotein (RNP)-based knock-in

For Cas12a RNP assembly, 0.5 μL of 100 μg/μL poly(L-glutamic acid) (PGA; 15– 50 kDa; Sigma-Aldrich) was combined with 0.48 μL *TRAC*-specific CRISPR-RNA (crRNA) (IDT) and mixed thoroughly. Then, 0.4 μL of Alt-R A.s. Cas12a Ultra (IDT; 10 μg/μL) was added and incubated for 15 min at RT to form RNPs, followed by addition of 1 μL HDRT (1-2 μg/μL). The mixture was kept on ice until electroporation.

#### Electroporation

Cells were resuspended in 20 μL of ice-cold P3 electroporation buffer (Lonza) at a concentration of 1–1.5 × 10⁶ cells per reaction and immediately used for electroporation. The cell suspensions were mixed with the prepared reagent mix and transferred to a 16-well electroporation strip (Lonza). To ensure proper contact and eliminate air bubbles, strips were gently tapped on the bench before electroporation. Electroporation was performed on a 4D-Nucleofector (Lonza) using program EH-115. Immediately after, 100 μL of pre-warmed T cell medium was added per well. Cells were gently resuspended and transferred to 96-well round-bottom plates containing 150 μL pre-warmed T cell medium per well at a final density of 0.5 × 10⁶ cells/well.

#### HDR enhancing drugs

For experiments involving pharmacological HDR enhancement, T cells were transferred to T cell medium containing 0.5 μM HDR Enhancer v2 (IDT), AZD (MedChemExpress), Nedisertib (M3814)(Selleckchem) and optionally Novobiocin (MedChemExpress) or ART558 (GLPBIO) immediately after the recovery step post-electroporation. A medium change or initial cell split was performed within 24 hours post-electroporation using HDR enhancer-free T cell medium. Cells were subsequently expanded in 96-well round-bottom or 24-well plates and split every 2-3 days.

### Flow Cytometry

Flow cytometry was used to assess gene editing efficiency on day 4 post-electroporation and to evaluate the functionality of the generated CAR T cells. All measurements were performed on a CytoFLEX LX (Beckman Coulter) and analyzed using FlowJo v10 (BD).

Cells were transferred to 96-well U-bottom plates and washed with PBS (centrifuged at 400 g for 5 min at room temperature). A staining master mix containing fluorophore-conjugated antibodies was prepared in PBS and added at 20 μL per well. Cells were incubated for 15 min at 4 °C unless otherwise specified, followed by one or two PBS washes prior to acquisition. To prevent cross-reactivity from the anti-Fc antibody used for CAR detection, extracellular staining began with anti-Fc AF647 and a viability dye, followed by two washes before staining additional surface markers.

### Animal experiment

Experiments involving mice were performed at Baylor College of Medicine in compliance and with approval from the Institutional Animal Care and Use Committee (IACUC) under ID: AN-5551. In brief, immunodeficient mice were infused with 0.5 × 10^6^ Nalm-6 cells (expressing GFP and fireflyluciferase) via tail vein injection. Seven days later, 0.5 × 10^6^ CAR^+^ T cells were infused intravenously. *TRAC* KO T cells were injected at the same total number as the TCR-deficient *CD3ζ*-CAR T cells. T cells were cryo-preserved in FCS supplemented with 10% DMSO on day 14 post blood collection then thawed, washed and formulated in PBS 3 hours prior to application. Tumor burden was assessed using bioluminescence imaging. Mice were anesthetized with Isoflurane and received intraperitoneally 150 mg/kg D-Luciferin (Revvity) dissolved in water. BLI was performed with the In Vivo Imaging System (Lumina III, Revvity). Animal welfare, body weights and general health conditions were controlled. Mice were euthanized after individual evaluation for each mouse with body weigh loss >20% or ethical end points reached.

### Cytotoxicity Assay of Allo-Reactive T Cells Against Multiplexed Gene-Edited T Cells

To assess resistance of gene-edited T cells to alloreactive killing, allo-reactive T cells were preferentially expanded using allogenic feeder cells for allogeneic stimulation. CD8⁺ T cells were enriched by MACS and co-cultured 1:1 with irradiated (30 Grey), CD3-depleted PBMCs from a second donor on the day of isolation and again on day 7, followed by expansion. 1 day after stimulation with feeder cells, T cell medium containing cytokines (IL-7 and IL-15) was added for cell expansion.

Cytotoxicity assays were performed using target T cells (mock-electroporated or gene-edited lacking HLA class-I and -II expression) from the same donor used for allogeneic stimulation. 25,000 CFSE (Thermo Fisher Scientific) labeled target cells were seeded per well in 96-well round-bottom plates. Allo-reactive effector T cells were added at varying effector-to-target (E:T) ratios (2:1 to 0.5:1) and wells containing only target cells served as controls. Plates were briefly centrifuged (100 g, 1 min, RT) and incubated for 16 h at 37 °C and 5% CO₂. For the readout, 100 μL of cell suspension was mixed with 100 μL of DAPI in PBS (1:10,000), incubated at 4 °C for ≥10 min, and 30 μL were acquired on a flow cytometer. Cytotoxicity was quantified based on the number of viable CFSE^+^ target cells compared to target-only controls.

### Nalm-6 co-culture assay

CAR-T cells tumor control was assessed in vitro via live cell imaging of GFP- expressing Nalm-6 tumor cells on an Incucyte device (Sartorius) in phenol red-free RPMI 1640 media (Gibco). CAR-T cells were expanded for 14 days following electroporation and subsequently rested for 24 hours in cytokine-free medium prior to the assay. A total of 2 x 10^4^ GFP-labeled Nalm-6 cells were seeded into a flat-bottom 96-well plates and co-cultured in a 10:1 target-to-effector cell ratio with T cells. To account for differences in initial editing efficiencies, all effector samples were adjusted to the same frequency of CAR^+^ T cells with *TRAC*-KO cells, ensuring an equal total cell number across conditions. The plates were then placed into the Incucyte device and a repeat scanning was performed over 48 hours with scans taken every 2 hours, capturing three images per well.

### Sanger Sequencing and Gene Editing Quantification

To quantify gene editing efficiency at the DNA level, genomic DNA (gDNA) was isolated on day 14 post-electroporation using the DNeasy Blood and Tissue Kit (Qiagen). Target regions at the *RASA2* and *FKBP12* loci were amplified via PCR using KAPA HiFi polymerase (Roche) with primers designed in COSMID^67^ to generate 500– 700 bp amplicons. Primer specificity was confirmed in silico using Primer-BLAST^68^ and validated by electrophoresis on a 1.5% agarose gel. PCR products were purified using the DNA Clean & Concentrator-5 Kit (Zymo Research) and submitted for Sanger sequencing (LGC Genomics).

Base editing efficiency was quantified using EditR^69^ and indel frequencies were estimated using ICE analysis^70^ (Synthego). Genomic DNA from mock-electroporated cells of the same donors served as negative controls for both analyses.

### CAST-seq

Genomic DNA was extracted using the NucleoSpin Tissue kit (Macherey-Nagel). CAST-Seq analyses were performed following the previously described protocol^71^, with some adjustments to the workflow^72^. In brief, the average fragmentation size of the genomic DNA was aimed at a length of 500 bp. The libraries were sequenced on a NovaSeq 6000 using 2 × 150 bp paired-end sequencing (GENEWIZ, Azenta Life Sciences). Changes were made to the bioinformatic pipeline to enhance specificity. Sites under investigation were categorized as OMT if the p value met the cutoff of 0.005. In addition, further features were incorporated in the CAST-Seq algorithm, including barcode hopping annotation and an update to the coverage analysis to reduce the execution time by aligning the gRNAs only to the most covered regions for each site. Two technical replicates from samples of two different donors were used for the CAST-Seq analysis. Only sites identified as significant in at least 3 out of the total 4 replicates were considered as putative events.

### Digital droplet polymerase chain reaction (ddPCR)

To quantify chromosomal translocations, ddPCR assays were designed to detect balanced translocations involving the *TRAC, RASA2*, or *FKBP12* loci (**Suppl. Table 3**). The RPP30 gene served as a reference control, as previously described.

For each reaction, 75 ng of genomic DNA digested with HindIII-HF (NEB, Germany) was used as the template in a 22 μL ddPCR mix containing:

- 5.5 μL of 4x dPCR Multiplex Supermix (Bio-Rad)
- 1.25 μL (10 μM) of each forward and reverse primer for the reference gene
- 1.25 μL (5 μM) of FAM-labeled probe for the reference gene
- 1.25 μL (10 μM) of each forward and reverse primer for the target gene
- 1.25 μL (5 μM) of HEX-labeled probe for the target gene
- Nuclease-free water to final volume

Droplet generation was carried out using the QX200 Droplet Generator (Bio-Rad) by loading 20 μL of sample mix and 70 μL of droplet generation oil into a DG8™ Cartridge, sealed with a DG8™ Gasket. Empty wells were filled with ddPCR Buffer Control.

Following droplet generation, droplets were carefully transferred into a 96-well PCR plate, sealed with pierceable foil using the PX1™ PCR Plate Sealer at 185 °C for 5 s. PCR was performed according to the manufacturer’s cycling protocol. Samples generating fewer than 15,000 accepted droplets or more than 99% positive droplets for the reference gene (*RPP30*) were excluded and repeated.

Primers and probes were synthesized by Integrated DNA Technologies (IDT). All other reagents and equipment were sourced from Bio-Rad.

### Data Analysis, visualization and text editing

Experimental data were collected in Excel and analyzed using GraphPad Prism 9, with statistical methods detailed in the main text. Figures and schematics were created in BioRender.com. Text clarity and grammar were improved using ChatGPT (OpenAI)

## Declarations

### Ethics approval and consent to participate

The study was performed in accordance with the Declaration of Helsinki. Peripheral blood from healthy human donors was obtained after informed and written consent (Charité ethics committee approval EA4/091/19 and EA1/052/22, Baylor College of Medicine IRB approval H-45017).

### Availability of data and materials

The ABE8.20m and BE4 were gifts from Nicole Gaudelli (Addgene plasmid #136300 and #138340). The original HDR templates for *TRAC* and *CD3*ζ were deposited with Addgene (Addgene plasmids #215769 and #215759). To encourage broad adoption of BEKI, the BEKI-adapted HDR template plasmids for CD19-CAR knock-in at *TRAC* and *CD3ζ* (*CD247*) have also been deposited with Addgene (Addgene plasmids #247056 and #247057). The IVT plasmid backbone contains proprietary 5’-UTR and 3’-UTR sequences and was procured from TriLink Inc. under a non-disclosure agreement. All other construct sequences can be found in **Suppl. Table 1.**

### Author contributions

VG designed the study, planned and performed experiments, analyzed results, interpreted the data, and wrote the manuscript. L.J.B., C.F.-G., L.H., L.M.H, A.-M.N., Y.P., C.L.F., I.K. and M.P. performed experiments, analyzed results, interpreted data and edited the manuscript. M.S., R.K. and S.S. performed experiments, analyzed results and interpreted data. G.A. analyzed and interpreted CAST-seq data. T.Ca. supervised CAST-Seq analysis, provided reagents, interpreted data, and edited the manuscript. H.-D.V and PR provided reagents, interpreted data, and edited the manuscript. J.K. and D.L.W. designed and led the study, planned experiments, interpreted data, and wrote the manuscript. All authors discussed, commented on, and approved the manuscript in its final form.

## Acknowledgements

The authors would like to thank Cliona Rooney and Lisa Rollins (both, Baylor College of Medicine) for their help in the animal studies. The project received funding from the European Union under grant agreement no. 101057438 (geneTIGA: genetiga-horizon.eu) to T.Ca., H.-D.V., P.R. and D.L.W. Views and opinions expressed are those of the author(s) only and do not necessarily reflect those of the European Union or the European Health and Digital Executive Agency (HADEA). Neither the European Union nor the granting authority can be held responsible for them.

V.G., J.K. and D.L.W. were supported by the SPARK-BIH Program by the Berlin Institute of Health, Germany. We also acknowledge funding from the German Federal Ministry of Education and Research (BMBF) within the Medical Informatics Funding Scheme EkoEstMed–FKZ 01ZZ2015 (G.A.).

## Supplementary Figures

**Suppl. Fig 1.**
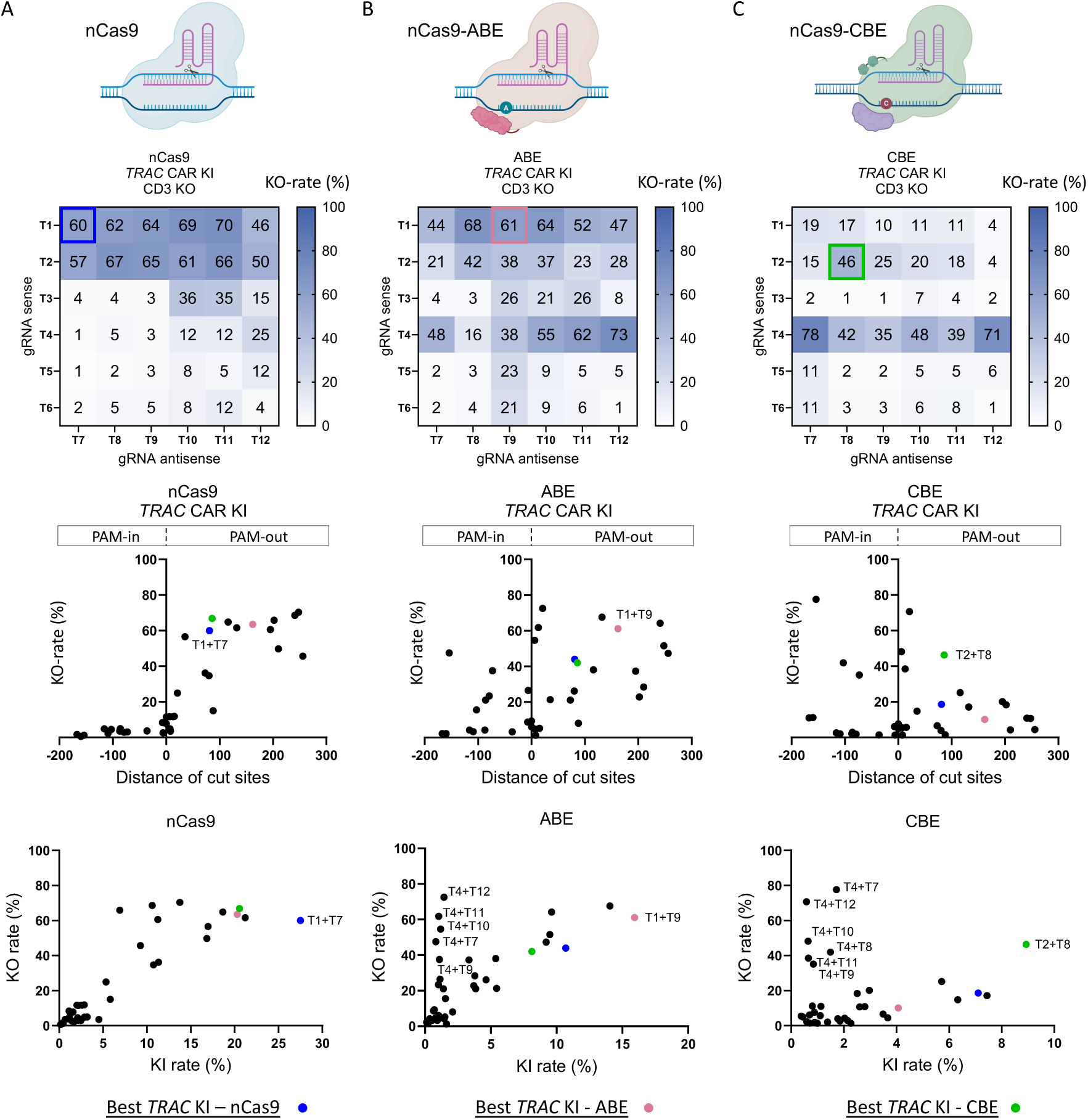
KO-efficiency when performing double nick-mediated *TRAC* editing using nCas9, ABE or CBE. (**A**) KO-efficiency using a D10A nCas9 was evaluated on day 4 after electroporation via flow cytometry (n=2 healthy donors). The results are displayed as a heatmap corresponding to Fig. 1B) and as a graphs plotting the KO-rate against the Nick distance or KI-rate. (**B**) KO-efficiency using a ABE8.20-m was evaluated on day 4 after electroporation via flow cytometry (n=2 healthy donors). The results are displayed as a heatmap corresponding to Fig. 1B) and as a graphs plotting the KO-rate against the Nick distance or KI-rate. (**C**) KO-efficiency using a CBE (BE4) was evaluated on day 4 after electroporation via flow cytometry (n=2 healthy donors). The results are displayed as a heatmap corresponding to Fig. 1B) and as a graphs plotting the KO-rate against the Nick distance or KI-rate.

**Suppl. Fig 2.**
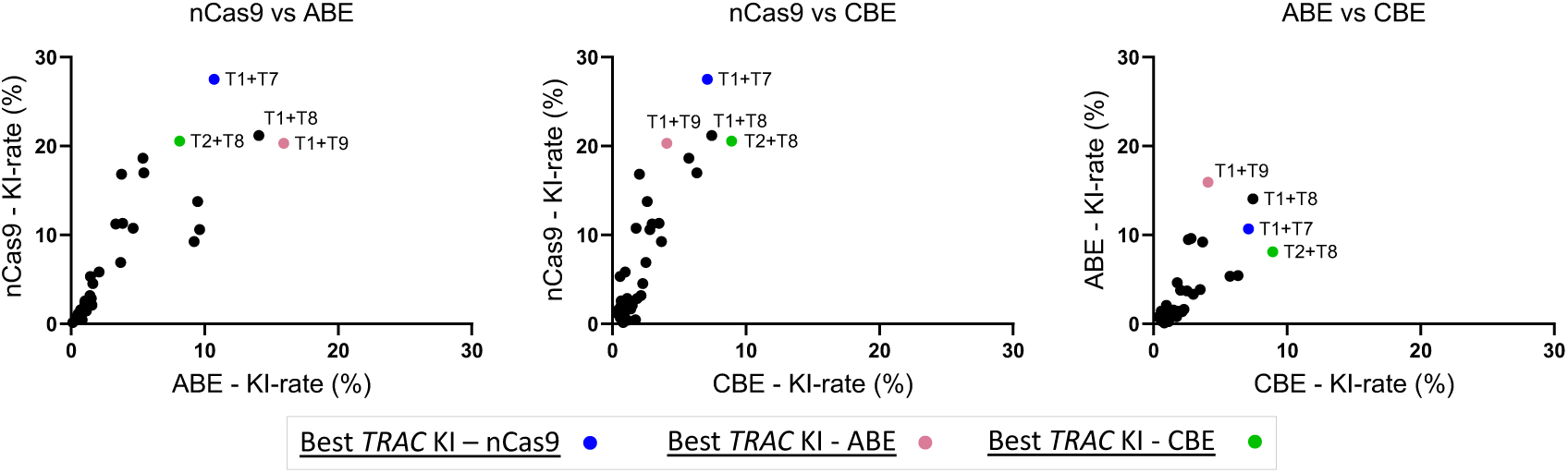
Correlation of KI-efficiency when using different editors. **Left:** The mean KI-efficiency using a D10A nCas9 was plotted against the mean KI-efficiency using ABE (n = 2 healthy donors). **Middle:** The mean KI-efficiency using a D10A nCas9 was plotted against the mean KI-efficiency using CBE (n=2 healthy donors). **Right:** The mean KI-efficiency using an ABE was plotted against the mean KI-efficiency using CBE (n=2 healthy donors).

**Suppl. Fig 3.**
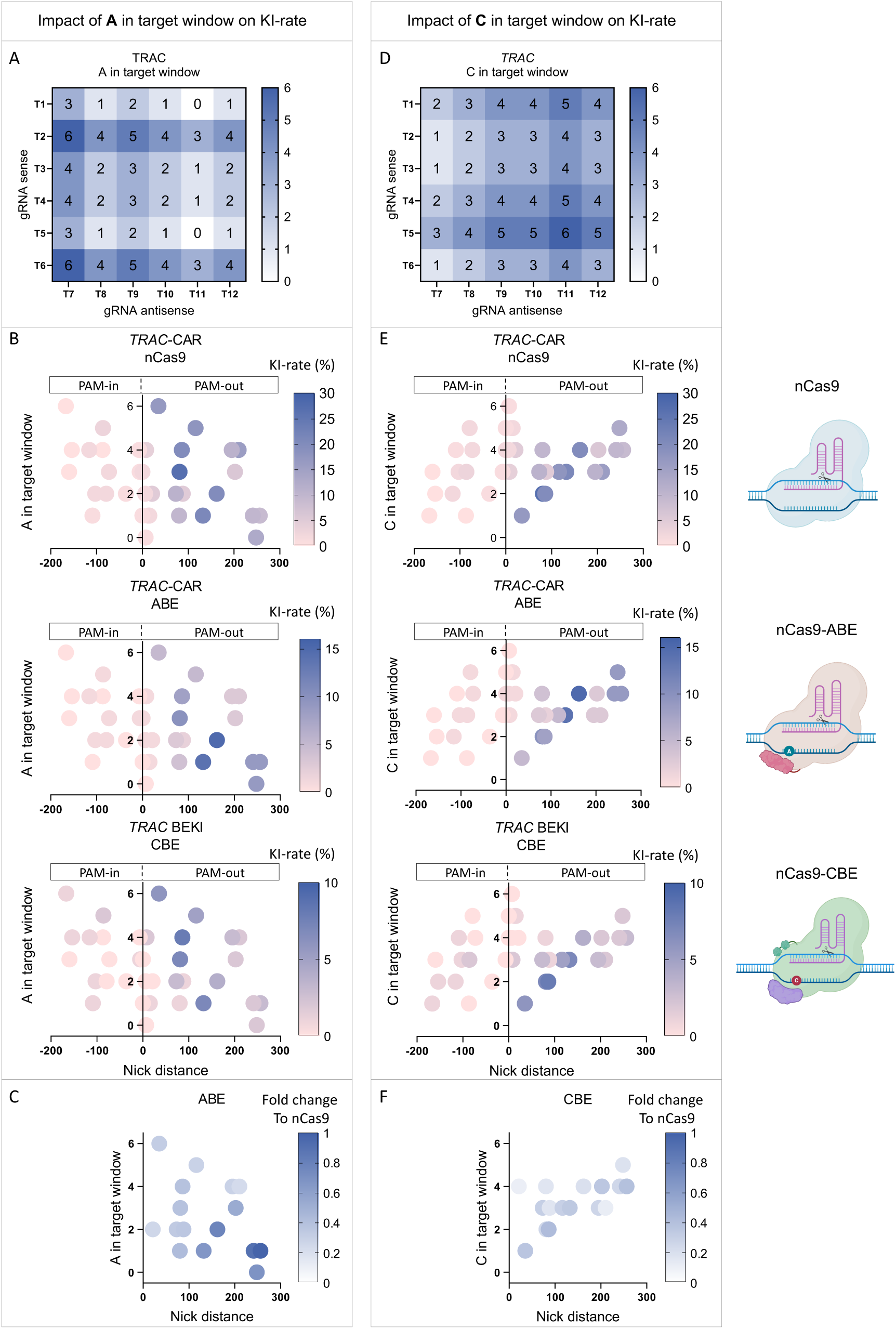
Impact of potential base edits on BEKI-efficiency using ABE or CBE. (**A**) Number of adenines (A) in the editing windows of each sgRNA pair for *TRAC* editing used in Fig.1. (**B**) The KI-rate (color intensity) using nCas9, ABE or CBE was plotted against the number of A in the editing window and the Nick distance (mean of: n=2 healthy donors). (**C**) Fold change of the KI-rate of ABE compared to nCas9 editing using sgRNA pairs in a PAM-out orientation. (**D**) Number of cytosines (C) in the editing windows of each sgRNA pair for *TRAC* editing used in Fig.1. (**E**) The KI-rate (colour intensity) using nCas9, ABE or CBE was plotted against the number of C in the editing window and the Nick distance (mean of: n=2 healthy donors). (**F**) Fold change of the KI-rate of CBE compared to nCas9 editing using sgRNA pairs in a PAM-out orientation.

**Suppl. Fig 4.**
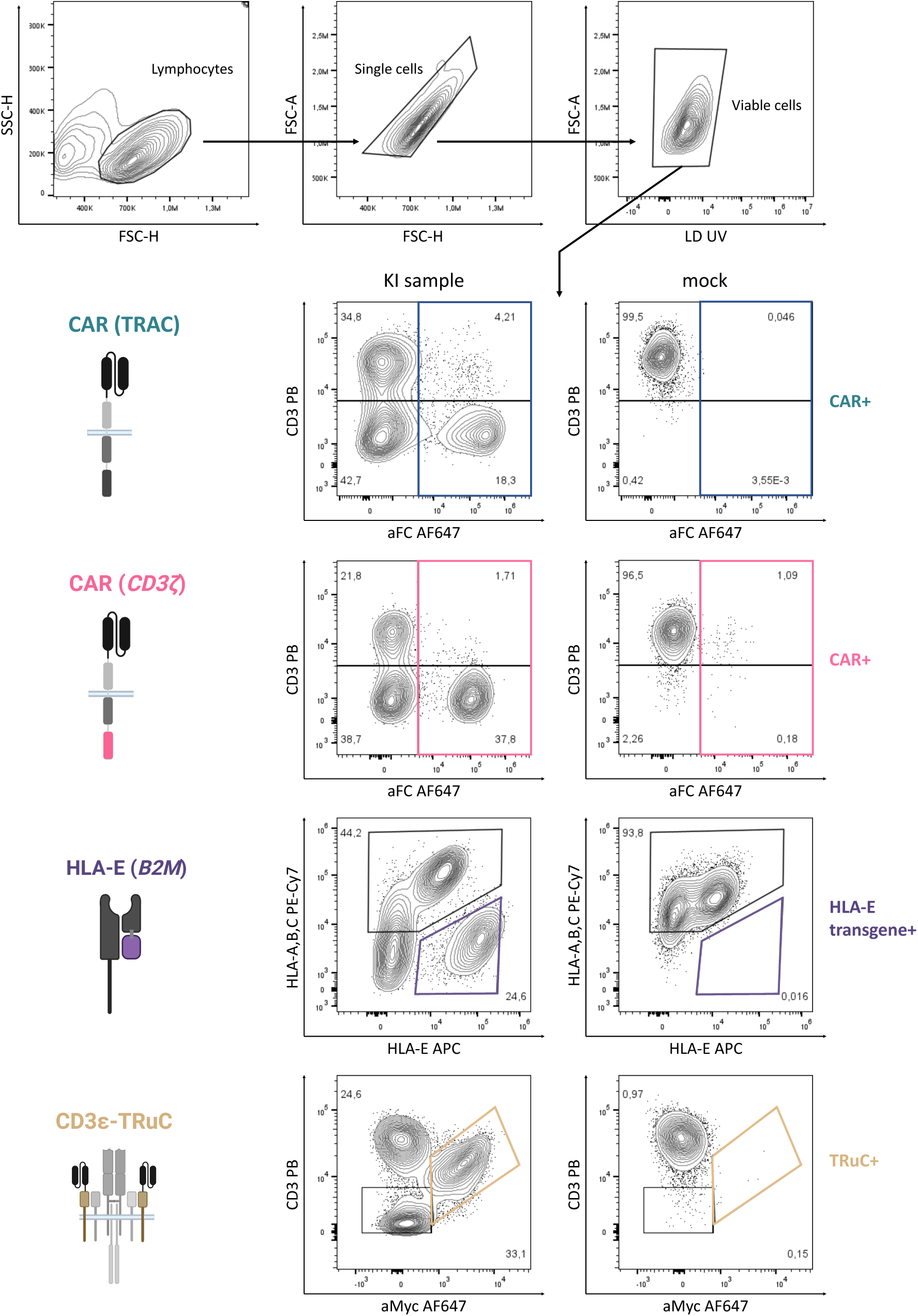
Gating strategies for flow cytometric analysis of the KI-efficiency. Exemplary flow plots are shown for the KI of a CAR into *TRAC*, a truncCAR into *CD3*ζ, HLA-E into *B2M* or a TRuC into *CD3*ε (all generated using BEKI with DNA-PKi + Polθi).

**Suppl. Fig 5.**
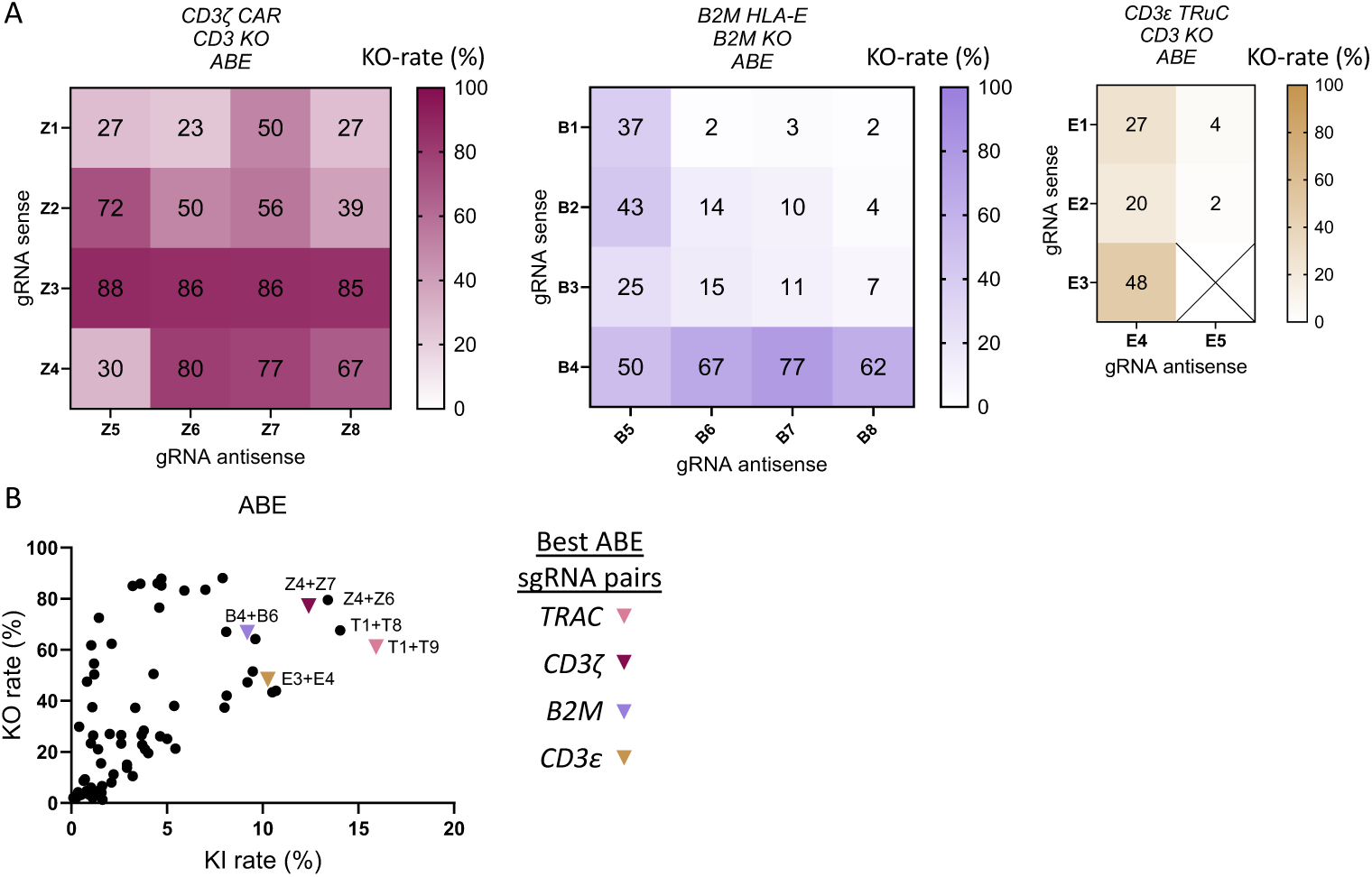
KO-efficiency of BEKI at the *CD3ζ,, B2M* or *CD3ε* locus using ABE. (**A**) KO-efficiency using a ABE8.20-m was evaluated on day 4 after electroporation via flow cytometry (n = 2-4 healthy donors). The results are displayed as a heatmap corresponding to Fig. 2. (**B**) The KI-rate of BEKI-edited cells at the *TRAC, CD3ζ,, B2M* or *CD3*ε was plotted against the respective KO-rates. The best sgRNA combinations for the 4 loci are highlighted.

**Suppl. Fig 6.**
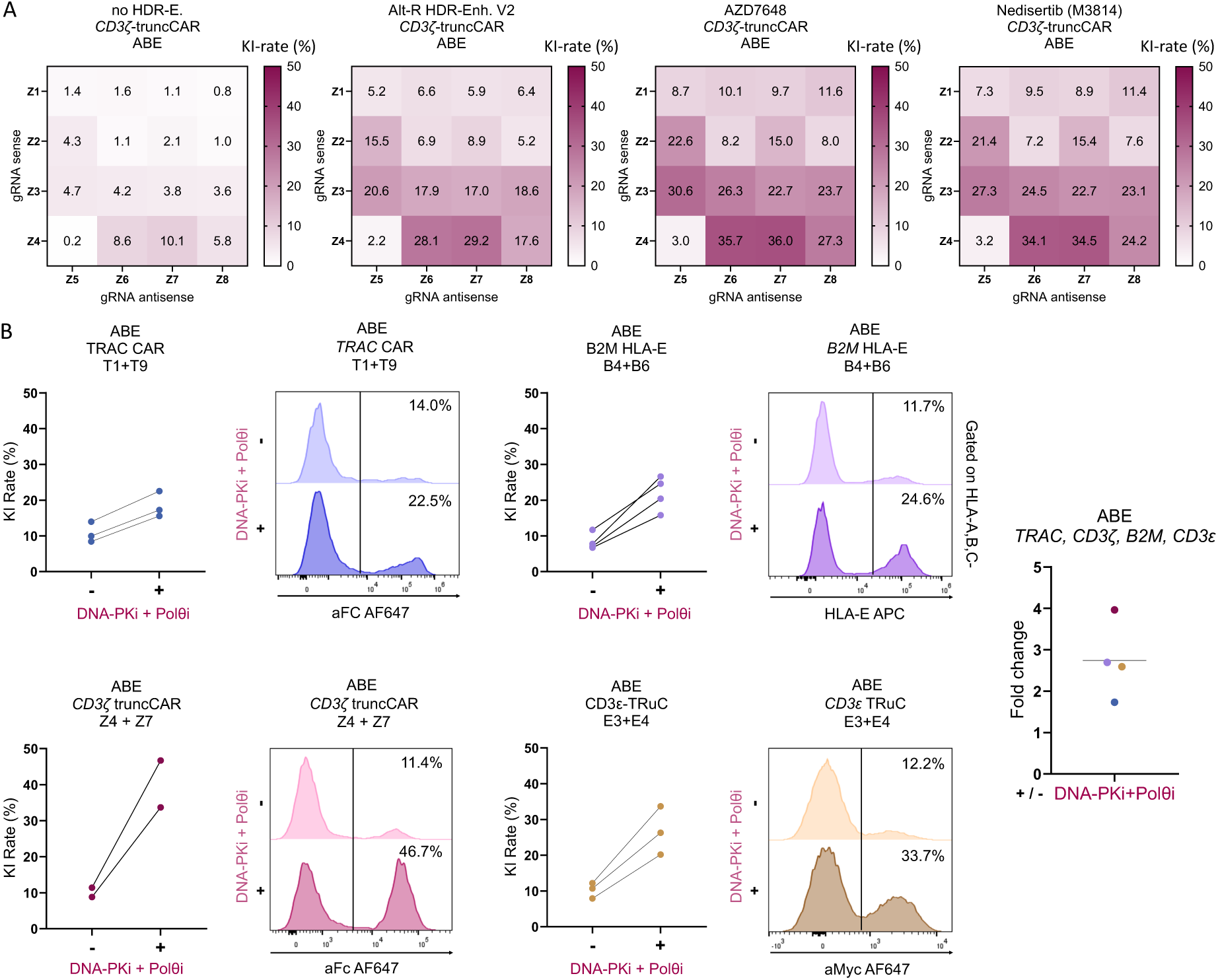
BEKI gene editing-efficiency at multiple therapeutically relevant loci can be enhanced using small molecule inhibitors. (**A**) BEKI of a truncCD19 CAR into the *CD3*ζ locus w/o or with different DNA-PK inhibitors (Alt-R HDR enhancer V2, AZD7648, Nedisertib (M3814)) for the sgRNA combinations shown in Fig.2 (n=2 healthy donors). (**B**) KI-rate of the most efficient combination for ABE-mediated KI into the *TRAC, CD3*ζ*, B2M* or *CD3*ε locus (n=2-4 healthy donors). Representative flow cytometry histograms show editing outcomes when adding DNA PK and Polθ inhibitors.

**Suppl. Fig. 7.**
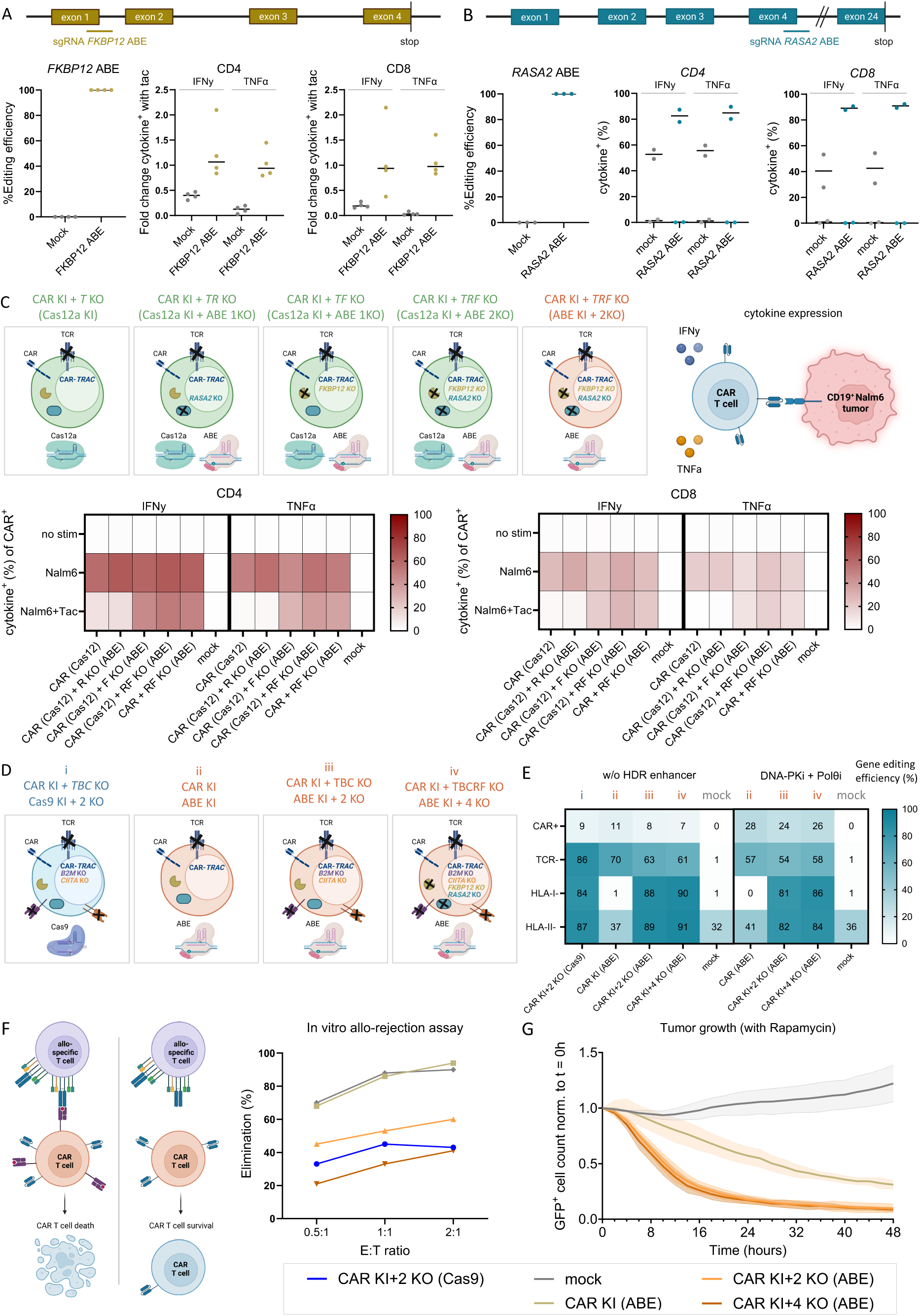
Generation of multiplex-edited CAR T cells with base editing-mediated splice site disruption of *FKBP12* and *RASA2*. (**A**) BE-efficiency of *FKBP12* was assessed using sanger sequencing followed by quantification of the base editing frequency via EditR. The mock electroporated or base edited cells were stimulated using aCD3/aCD28 antibodies with or w/o tacrolimus (Tac, final concentration of 6 ng/mL) before flow cytometric analysis of the TNFα and IFNγ expression of CD4 and CD8 T cells. The fold change of expressing cytokines in the presence of Tac was calculated compared to the frequency w/o Tac treatment. (**B**) BE-efficiency of *RASA2* was assessed using sanger sequencing followed by quantification of the base editing frequency via EditR. The mock electroporated or base edited cells were stimulated using aCD3/aCD28 antibodies with or w/o tacrolimus (Tac, final concentration of 6 ng/mL) before flow cytometric analysis of the TNFα and IFNγ expression of CD4 and CD8 T cells . Unstimulated cells are included as a control (lighter color). (**C**) Gene edited CD19 CAR T cells or mock electroporated T cells were stimulated using CD19+ Nalm 6 tumor cells in the presence or absence of Tac. Flow cytometric analysis of the TNFα and IFNγ expression of CD4 and CD8 T cells was performed after 6 hours. (**D**) Schematic overview of the different gene editing strategies targeting the *TRAC, RASA2,FKBP12, CIITA and B2M* genes. (**E**) Gene editing efficiency of CAR KI and *TRAC* (CD3), *B2M* (HLA-A,B,C) and *CIITA* (HLA-DR,DP,DQ) KO was determined by flow cytometry (n=3 healthy donors). (**F**) Allo-specific CD8+ T cells were generated against another donor by adding irradiated CD3-cells twice. The allo-specific cells were co-cultured with CFSE-labled mock electroporated or gene edited cells from the donor used for stimulation at effector to target ratios of 0.5:1, 1:1 and 2:1 (n=1 healthy donor). The number of CFSE+ cells was determined and the elimination frequency quantified using wells containing only target cells. (G) Tumor growth of the GFP^+^ Nalm6 cell line in a 10:1 co-culture with BEKI CAR-T cells in the presence of rapamycin (100 nM) was monitored over 48 hours using the Incucyte live-cell analysis system (n = 2 healthy donors in 3 techn. replicates each).

**Suppl. Fig 8.**
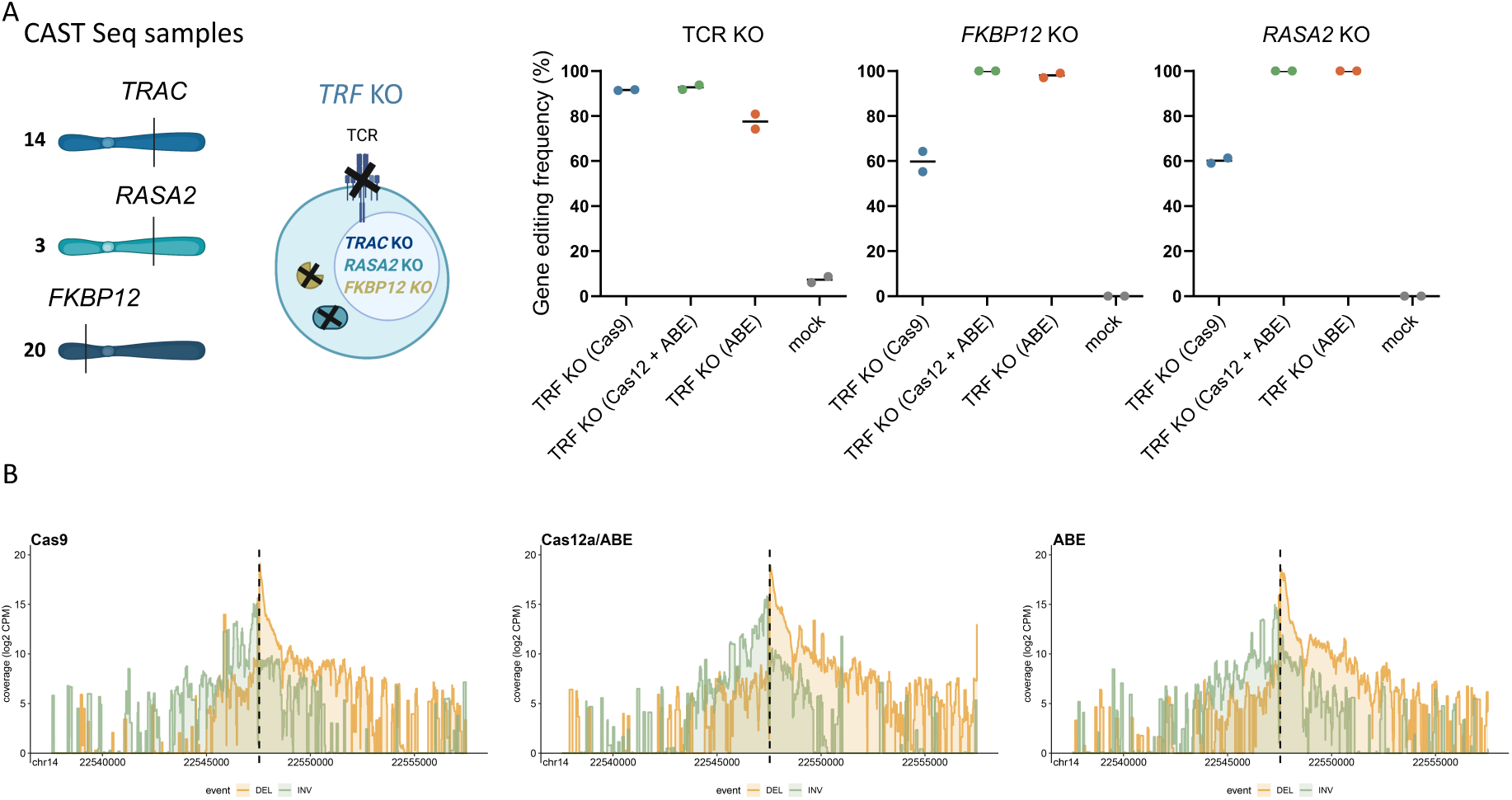
Gene editing efficiency of *TRAC, RASA2* and *FKBP12*-edited samples used for CAST-Seq analysis. (A) Gene editing efficiency of the TRAC KO was determined by flow cytometry. KO- or BE-efficiency of *RASA2* and *FKBP12* gene were assessed using sanger sequencing followed by indel analysis using TIDE or base editing frequency via EditR (n=2 healthy donors). (B) Deletions (yellow) and inversions (green) at the *TRAC* locus detected in cells edited with Cas9, Cas12a + ABE or ABE using CAST-Seq (excluding an HDR donor template).

